# Soil microbial diversity alters soil microhydrology through extracellular polymeric substance production

**DOI:** 10.64898/2026.06.03.729803

**Authors:** Yvonne Kan, Mariana Acevedo, Hailey Buell, Emily Herrera, Avery Swanton, Alonso Favela

## Abstract

Soil microbial communities have a variety of mechanisms to deal with emerging drought stress. One well-documented mechanism is increased microbial production of extracellular polymeric substances (EPS), which can potentially change the soil density and water holding capacity. Yet little is known about how microbial diversity influences the functional capacity of EPS formation and the resulting outcomes in water dynamics. To understand more about communal microbiome EPS production, we set up sterile mesocosms where we examined the effects of microbial diversity (high or low treatments) and nutrient input (supplement or deficient treatments) on these processes. To capture the microhydrology of the mesocosms, we measured water holding (WH), infiltration, evaporation, and soil properties we believe microbes are altering (EPS, soil aggregation). Our hypothesis stated that if diversity was artificially manipulated, then soil-water properties will be altered via production of EPS. We predicted that low diversity systems would have lower functional diversity, leading to less EPS production, moisture storage, and minimal changes from inert soil media. As predicted, we found that the high-diversity systems had a higher water retention and lower rates of water loss over time than low-diversity systems. This trend was magnified in the nutrient-supplemented treatment, suggesting that EPS production and subsequent water-holding traits are emergent features of the microbiome. Unexpectedly, we observed a correlation between the amount of water retained and the quantity of lipid EPS produced. This suggests that EPS composition, rather than quantity, is determinative of a biofilm’s function. In conclusion, it appears that microbial diversity influences soil properties that are important to moisture retention within these systems. To date, the role that microbes and their diversity play in soil hydrology has been severely understudied, so this work aims to build ecological understandings of these systems. These findings are valuable, for if we learn how microbes manipulate soil moisture, we can apply these functions to advance sustainable agricultural practices and enhance ecosystem resilience to water scarcity in arid regions.

**Open Research Statement:** Upon publication data, and code will be made available through Zenodo. Sequencing data will be uploaded to NCBI SRA.

## INTRODUCTION

Drought has emerged as one of the most pervasive and consequential climatic stressors affecting both human and natural ecosystems (Seo et al., 2025). Unlike acute disturbances, drought develops through sustained precipitation deficits that progressively deplete terrestrial water storage, particularly soil moisture. In fact, recent global analyses indicate that terrestrial water storage has declined at nearly twice the rate of Greenland ice mass loss during the early twenty-first century (Seo et al., 2025). These trends reflect a fundamental shift in the global water cycle, driven by declining precipitation frequency and rising soil temperatures, with far-reaching consequences for managed and natural systems alike.

In agriculture, widespread drought is projected to account for 30–90% of global crop yield losses (Dietz et al., 2021), intensifying dependence on supplemental irrigation and placing growing pressure on already-strained freshwater resources. Beyond agriculture, persistent drought can reduce plant community productivity by up to 40% (Smith et al., 2024), with cascading consequences for animal communities and ecosystem stability. Together, these impacts underscore an urgent need to identify mechanisms that enhance soil water retention and buffer ecosystems against hydrological stress.

Over the past decade, soil microbial communities have been increasingly recognized as key regulators of soil habitat structure, influencing fertility, crop productivity, and plant tolerance to both biotic and abiotic stressors (Iqbal et al., 2025). Microorganisms comprising these soil communities are able to modify their physical and chemical environment through a wide range of biochemical and biophysical processes (Philippot et al., 2024). Yet, despite growing interest in microbiome-based solutions to climate change, our understanding of how to harness microbial functions to stabilize soil water dynamics remains limited. In response to drought, soil microorganisms employ diverse survival strategies, including the formation of protective biofilms: dense, multi-species, syntrophic assemblages thought to play a central role in moisture retention (Costa et al., 2018). These biofilms are constructed from extracellular polymeric substances (EPS), a heterogeneous matrix composed of polysaccharides, proteins, extracellular DNA, and lipids.

Understanding how EPS contributes to soil hydrology therefore requires characterization, not just of individual EPS producers, but of the community-level processes that govern EPS accumulation and composition. EPS composition and function are highly variable and taxon-dependent, with distinct microbial lineages producing polymers that differ in chemical structure, hydrophilicity, and mechanical stability (Schmid et al., 2015). Importantly, EPS production might not solely be a property of individual taxa, but an emergent feature of microbial communities shaped by interspecific interactions and metabolic cross-feeding (Xavier and Foster, 2007). Despite this complexity, the role of microbial diversity in governing EPS formation and its consequences for soil hydrology remains poorly resolved.

Most current insights into EPS function in soils derive from single-isolate studies or microbe-independent experimental systems. For example, Pérez et al. (2022) demonstrated that inoculation with a single high-EPS-producing bacterium could alter soil hydrodynamics in sandy substrates by modifying pore-scale physical and chemical properties. Similarly, Volk et al. (2016) showed that direct addition of EPS from a *Pseudomonas* culture reduced saturated hydraulic conductivity in soil matrices. While these studies provide compelling mechanistic evidence that EPS can influence soil physical structure, they do not capture the ecological complexity of intact microbial communities. Observational surveys further suggest that soil EPS pools are sensitive to land-use change, with measurable consequences for soil physical properties (Redmile-Gordon et al., 2020). However, the mechanisms by which EPS mediates soil water retention remain unresolved (Redmile-Gordon et al., 2014), and we lack experimental tests of how microbial diversity and environmental context, particularly nutrient availability, interact to govern EPS accumulation and its hydrological consequences. This knowledge gap limits our ability to deploy microbiome-mediated processes to manage critical ecosystem functions such as plant health, soil moisture retention, and hydrological regulation, despite clear indications that microbial EPS could play a transformative role. Given these areas for discovery, we hypothesize that soil microbial diversity will be an important driver of EPS pools within a soil system, and that alterations to soil diversity can alter the hydrology of a soil.

Here, we experimentally test whether microbial biodiversity contributes to EPS formation in soils and whether this, in turn, shapes soil hydrological properties. We developed controlled soil mesocosms in which we could manipulate microbial diversity and nutrient pools while enabling precise quantification of soil water storage, evaporation, and infiltration. Specifically, we asked: (i) whether high- and low-diversity microbial communities differ in their effects on soil hydrology; (ii) whether EPS accumulation and composition mediate the observed hydrological differences; and (iii) whether specific microbial taxa are tightly coupled to EPS concentrations and soil moisture retention. Together, our study provides experimental evidence linking microbial diversity to EPS formation and soil hydrological function, advancing a mechanistic framework for understanding how microbiomes may buffer soils against increasing drought stress.

## MATERIALS AND METHODS

The goal of this study was to understand how the microbiome and EPS production shape critical soil characteristics related to the movement of water. To examine this, we developed soil mesocosm systems that allowed us to measure bulk hydrological processes (such as infiltration and retention) and manipulate microbial composition in a controlled manner (**Figure 1**). All mesocosms were initialized with the same starting sterilized soil media, providing an equivalent baseline for soil properties across treatments. We manipulated two factors at the start of the experiment: the microbial composition inoculated into each mesocosm, and whether microbes were introduced with a nutrient or water solution. Hydrological measurements were recorded daily. At the conclusion of the experiment, EPS content, soil physical and chemical properties, and microbial community composition were quantified within each mesocosm.

**Figure 1.**
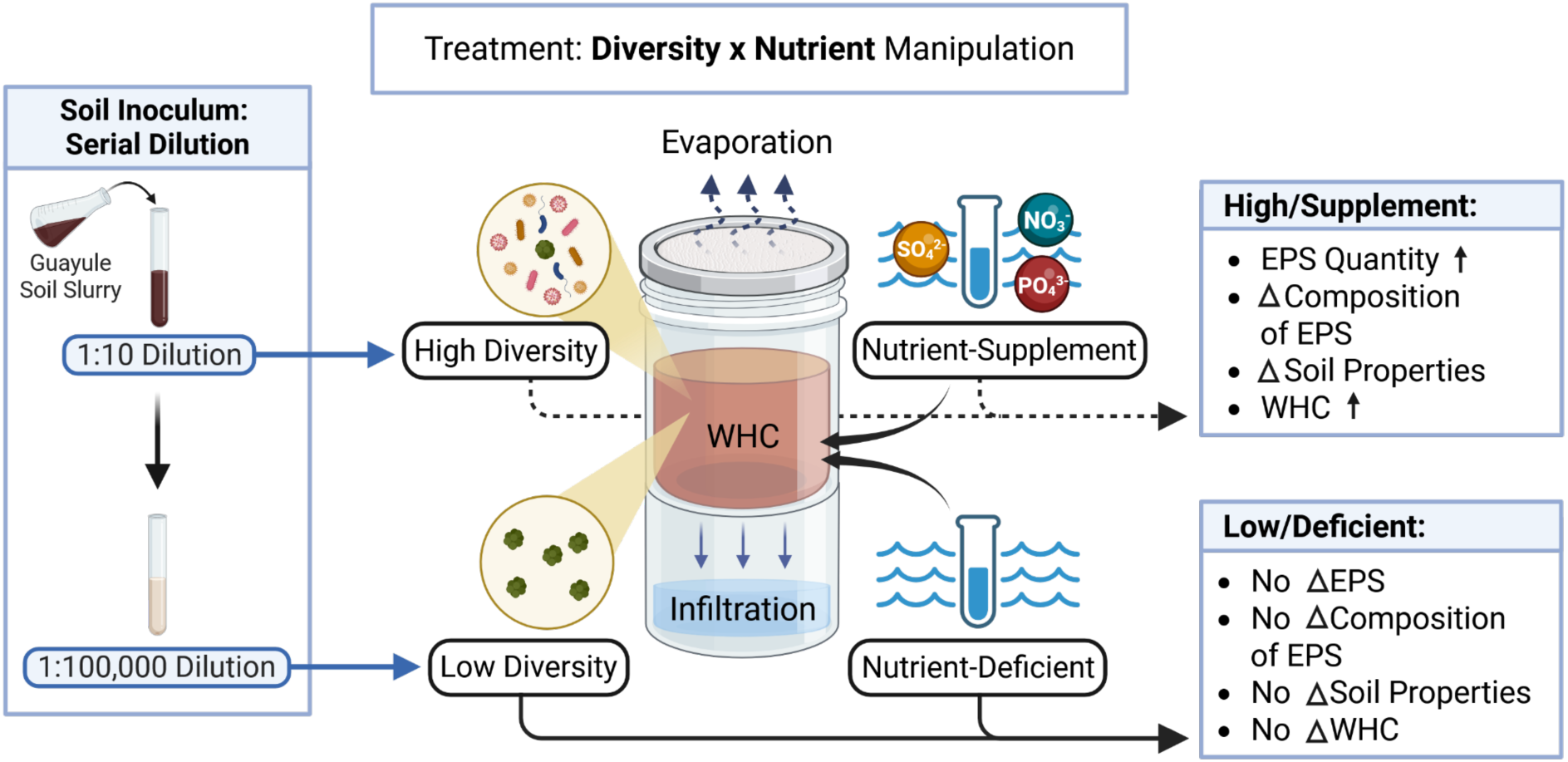
Methodology of soil mesocosms and development of diversity treatment alongside nutrient-availability treatment. The mesocosms were able to measure water-holding capacity, infiltration, and evaporation over time by tracking mass of water. Diversity and nutrient-availability within the system was manipulated by serial dilution and addition/exclusion respectively. Although we included many combinations of diversity levels and nutrient input, the figure shows the two treatment combinations resulting in the most significant difference in treatment effects. Created in BioRender. Kan, Y. Favela, A. (2025) https://BioRender.com/8prv1nx

### Soil Mesocosm Systems

Soil mesocosm systems were developed in-house using two stacked Magenta boxes. A hole (diameter 0.5cm) was drilled in the base of the upper box and fitted with a filter paper (size 2.5um/grade 4) to allow flow-through drainage into the lower box. A cellulose acetate (pore size 0.2um) filter was secured over the top of the mesocosm to maintain sterility while permitting evaporation.

The sterile soil medium consisted of a 1:1 (by volume) mixture of sieved soil (3.35 mm mesh) and sand, which was autoclaved (121°C, 15psi, 30 min, gravity cycle). For the diversity treatment, live inoculum soil was collected from agricultural soil at a guayule farm in Eloy, Arizona. A soil slurry was prepared at a ratio of 1 g soil to 10 mL nanopure water, this was our high-diversity treatment (**Figure 1**). This slurry was then serially diluted (1:10), and the final (1:100,000) dilution was used as the low-diversity treatments. 16S amplicon sequencing was performed on the inocula to confirm that dilution successfully manipulated microbial diversity as intended. The high-diversity inoculum contained a broad range of microbes spanning multiple phyla, with the predominant phyla being *Proteobacteria, Firmicutes,* and *Actinobacteriota*, followed by *Crenarchaeota* and *Verrucomicrobiota*. The low-diversity inoculum was derived from the same soil community but was diluted to substantially reduce diversity; this treatment was dominated almost exclusively by *Actinobacteriota*.

### Experimental Spin-up

To allow the soil media to first fully hydrate prior to inoculation, we initially treated the mesocosms with a nutrient-availability treatment. Each mesocosm was wetted with 100mL of either Hoagland’s nutrient solution (nutrient-supplemented) or sterile water (nutrient-deficient) (**Figure 1**). After an hour of letting the mesocosm soil media hydrate and drain of liquid, they were inoculated with 1mL of the diversity treatment slurry.

We then wet the systems again with another 100mL of sterile water and measured the initial mass of the system to start the water infiltration experiment. We regularly (once a day, 5 days a week) measured the mass of the systems to calculate the amount of water lost over time. This first cycle of wetting spanned about one month and is an initial spin-up phase to let the microbial community equilibrate and reach a stable state. This also mitigates some of the potential differences in abundance of the microbial community, since the serial dilution may have also manipulated the quantity/size of the microbiome, so that diversity is the only contributing factor.

After the initial spin-up phase, we did a single round of rewetting (100mL sterile water) to simulate flooding/water input events and observe how soil hydrology is altered over time. During the following rewetting, we tracked the mass of the water retained and water infiltrated through the soil profile. Once the systems had dried completely, we would sample the soil media by taking 0.5g of soil from each corner of the top Magenta box (thus 2.0g of sampled soil in total). These samples were used for the EPS and soil chemistry measurements.

### EPS Extraction and Quantification

On the previously mentioned soil samples, cation exchange resin (CER) extraction was performed as described in Zhang et al. 2023 (similar protocol in Redmile-Gordon et al., 2014; Frølund et al., 1996), then we filtered the extracts using SnakeSkin dialysis tubing (22mm 12-14kDa) placed in DI water. Briefly, the CER method works by mixing the soil with a charged cation resin that sequesters common Ca²⁺ and Mg²⁺ ions, destabilizing the electrostatic bonds anchoring EPS to cell surfaces and causing it to dissociate into the surrounding solution. This method is generally considered gentle, resulting in minimal cell lysis. Filtration through a cellulose membrane was necessary to isolate EPS produced solely by microbial activity. The colorimetric methods used to quantify EPS were sensitive to the ions present in the nutrient solution added at the start of the experiment. Filtering the samples removed these background contaminants, retaining only the polymers relevant to EPS characterization. Although we potentially lost some EPS molecules smaller than the 12-14 kDa pore size, this step greatly minimized background contamination and improved the accuracy of downstream quantification.

From this EPS extract, we characterized three main components of EPS (polysaccharides, proteins, lipids). While carbohydrates are traditionally considered the primary structural component of EPS, recent work suggests that other fractions may also play important roles in EPS architecture (Karygianni et al., 2020). Polysaccharides were quantified using the phenol-sulfuric assay (DuBois et al., 1956). Proteins were quantified using the modified Lowry assay (Redmile-Gordon et al., 2013; Rojas and Pavón, 2005). Lipids were quantified using a sulfo-phosho-vanillin (SPV) assay (Knight et al., 1972; Zhong, 2016). Colorimetric measurements were performed on a Tecan Infinite 200 Pro plate reader, with absorbance measured at wavelengths specific to each protocol.

### Soil Chemical and Physical Properties

Nitrate, ammonium, and organic carbon were measured from soil samples collected at the end of the experiment. Nitrogenous compounds were extracted using a KCl extraction procedure. Nitrate was quantified colorimetrically using the vanadium reduction method (Soils Lab University of Illinois, 2023; Hendrix and Braman, 1995), while ammonium was quantified using the salicylate method (Soils Lab University of Illinois, 2023; Nelson, 1983). Colorimetric measurements were performed on a Tecan Infinite 200 Pro plate reader.

Soil organic carbon (SOC) was quantified using a combustion method (Miller, 2013). Soil samples were weighed to an initial mass of 4.5g. They were then combusted in a muffle furnace and weighed to find the final mass of the soil samples post combustion. Mass of soil organic carbon loss was calculated using initial and final mass.

Soil aggregation was assessed using a dry sieve method. Dried soil was passed sequentially through sieves of decreasing mesh size (3.35, 2.36, 0.25, and 0.18 mm), with material retained at each fraction collected and weighed. Total sample mass was recorded prior to sieving, and any mass lost during the process was calculated as a percentage. The proportion of soil retained at each sieve size was used as a proxy for aggregate size distribution.

### Microbiome Characterization

At the end of the experiment, a small fraction of soil was frozen and lyophilized. Lyophilized soil was extracted using the Qiagen DNeasy PowerSoil Pro Kit following the manufacturer’s instructions. Post-extraction DNA was quantified using PicoGreen (Thermo Fisher Scientific) on a Tecan Infinite 200 Pro plate reader. Purified DNA samples were submitted for amplicon sequencing at the University of Arizona Genetics Core (UAGC). Amplicon sequencing targeted bacterial and archaeal 16S ribosomal RNA genes using the forward primer (515F) [5′-GTGYCAGCMGCCGCGGTAA-3′] and reverse primer (806R) [5′-GGACTACNVGGGTWTCTAAT-3′]. Library preparation was carried out by the UAGC. Samples averaged approximately 65,000 reads per sample.

Raw bacterial and archaeal 16S rRNA sequences were processed using the “DADA2” pipeline (Callahan et al., 2016) to generate an amplicon sequence variant (ASV) table. Singleton, chloroplast, and mitochondrial ASVs were removed prior to statistical analysis. ASV taxonomic annotations were done using the SILVA 138.1 database (Chuvochina et al., 2026) within the “DADA2” pipeline. Microbiome data were visualized using a combination of base R (R Core Team, 2013), “ggplot2” (Wickham, 2016), and “phyloseq” (McMurdie and Holmes, 2013).

## Statistical Analysis

This study included both continuous and endpoint sampling. As described above, hydrological measurements were taken throughout the duration of the experiment, while soil physical properties, EPS concentrations, and microbial composition were measured at the end. To account for this, two statistical models were used: one incorporating a temporal day effect for continuous data (Infiltration or Water Retention ∼ Diversity × Nutrient Treatment + (1|Replicate) + (1|Day)), and one containing only treatment effects for endpoint data (Infiltration or Water Retention ∼ Diversity × Nutrient Treatment + (1|Replicate)). Data from the initial spin-up phase were excluded from all analyses; all hydrological results presented are from the second rewetting cycle. Univariate data were analyzed using linear mixed effects models implemented in the R package “lme4” (Bates et al., 2015). Multivariate community data were analyzed using the R package “vegan” (Oksanen et al., 2026). The effects of diversity and nutrient treatment on microbial community composition were assessed using permutational analysis of variance (PERMANOVA) via the “adonis” function in “vegan”. Cohen’s D was reported throughout the manuscript to report the effect size.

## RESULTS

### Water Retention and Infiltration

Over the course of the experiment, both diversity and nutrient treatments significantly altered water retention but not water infiltration within our soil mesocosms (**Figure 2A-B**).

**Figure 2.**
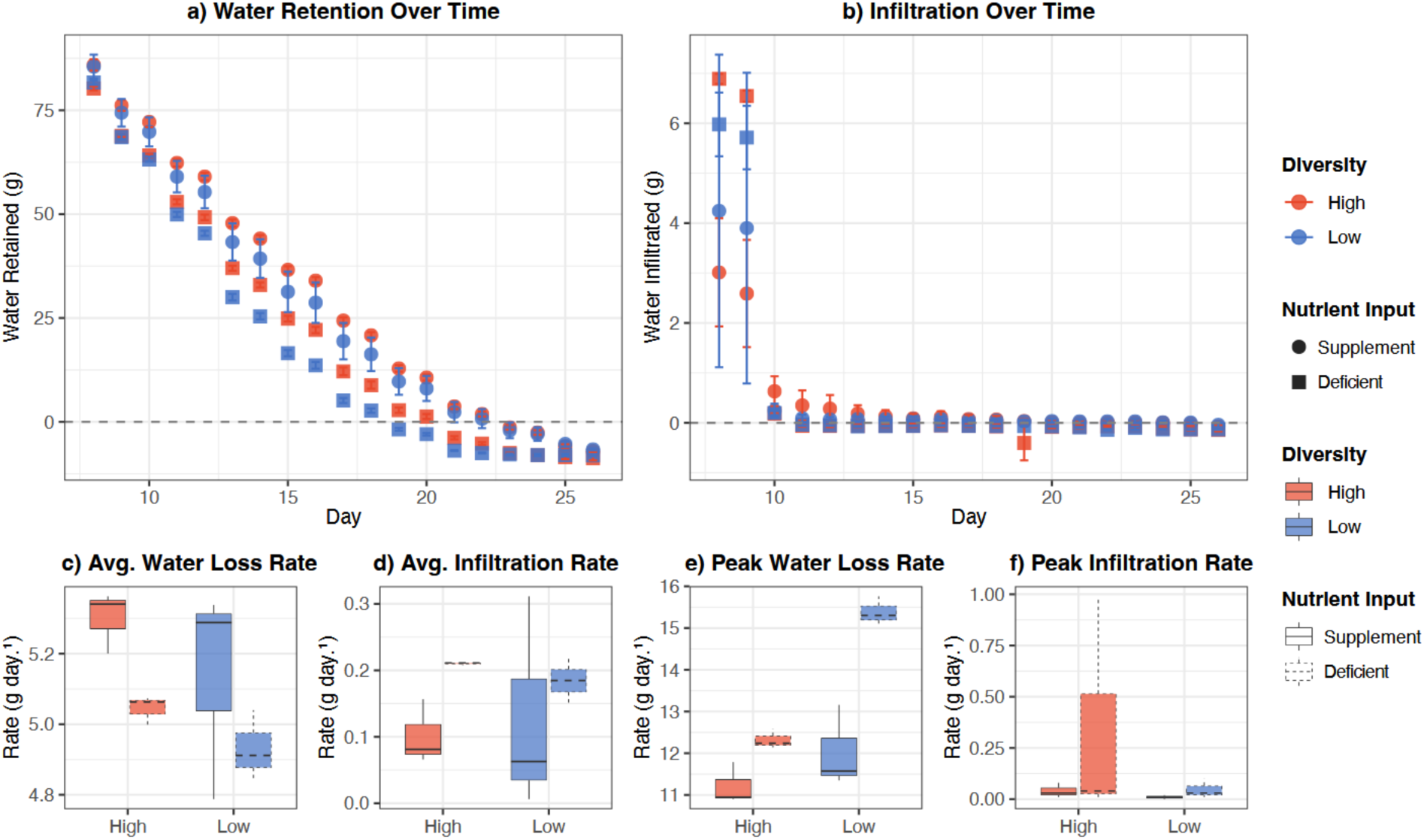
Hydrological dynamics of soil mesocosms under contrasting diversity and nutrient treatments. Mean (± SE) water retention (a) and infiltration (b) over time, with high-diversity mesocosms shown in red and low-diversity in blue; nutrient-supplemented and nutrient-deficient treatments are distinguished by circles and squares respectively. Average rates of water retention (c) and infiltration (d) were estimated as the linear slope of water volume over time. Peak rates of water loss (e) and peak infiltration (f) represent the maximum single-day change in water volume recorded per mesocosm. All rates are expressed in g day⁻¹; box plots show median, interquartile range, and 1.5× IQR whiskers; n = 3 mesocosms per treatment combination.

Nutrient treatment had the largest effect on water retention (d = -2.963, SE = 0.95, t = - 10.88, p < 0.001), followed by diversity (d = -1.156, SE = 0.95, t = -4.25, p < 0.001). Neither treatment significantly affected infiltration. The interaction between nutrient and diversity treatments was non-significant, suggesting these factors acted independently on water retention (p>0.05). The high-diversity nutrient-supplemented treatment retained the most water on average (30.3 ± 3.9 g), followed by low-diversity nutrient-supplemented (27.7 ± 3.9 g), high-diversity nutrient-deficient (21.9 ± 3.8 g), and low-diversity nutrient-deficient treatments (18.5 ± 3.9 g; Figure 2A).

At the onset of each rewetting cycle, water retention was similar across diversity treatments, with means differing by less than 1.41 g within nutrient treatments on Day 1. After 3–4 days, however, treatments began to diverge, with nutrient-deficient mesocosms losing water more rapidly than nutrient-supplemented ones (**Figure 2A**). Within nutrient-deficient systems, differences in water retention between high- and low-diversity treatments emerged by Day 4 and continued to widen until the mesocosms dried out completely, reaching a maximum difference of 8.5 g by Day 16. A similar but attenuated divergence was observed in nutrient-supplemented systems, where high- and low-diversity treatments differed by a maximum of 5.3 g, also peaking at Day 16.

Despite comparable average rates of water loss across treatments (ranging from -5.3 ± 0.09 g day⁻¹ in the high-diversity nutrient-supplemented treatment to -4.93 ± 0.10 g day⁻¹ in the low-diversity nutrient-deficient treatment), peak rates of water loss revealed greater treatment differentiation (**Figure 2C**). The low-diversity nutrient-deficient treatment exhibited the highest peak loss rate (−15.4 ± 0.35 g day⁻¹ at Day 13), followed by the high-diversity nutrient-deficient (−12.3 ± 0.24 g day⁻¹ at Day 12), low-diversity nutrient-supplemented (−12.0 ± 0.99 g day⁻¹ at Day 13), and high-diversity nutrient-supplemented treatments (−11.2 ± 0.50 g day⁻¹ at Day 13) (**Figure 2E**). Together, these results indicate that while mean loss rates were broadly similar, nutrient-deficient mesocosms, particularly low-diversity ones, experienced more intense peak water loss events.

Infiltration rates were low across all treatments throughout the second rewetting cycle, with average rates ranging from -0.10 ± 0.05 g day⁻¹ in the high-diversity nutrient-supplemented treatment to -0.21 ± 0.00 g day⁻¹ in the high-diversity nutrient-deficient treatment (**Figure 2D**). Peak infiltration rates similarly showed little treatment differentiation, except for the high-diversity nutrient-deficient treatment, which exhibited a notably higher peak infiltration rate (0.35 ± 0.56 g day⁻¹ at Day 21) compared to all other treatments, which peaked at or near zero (**Figure 2F**). The high variance in the high-diversity nutrient-deficient peak (SD = 0.56) suggests this may reflect an isolated infiltration event rather than a consistent treatment response.

### Extracellular Polymeric Substances

To investigate the potential biological mechanism underlying the observed diversity effects on water retention, we quantified microbial extracellular polymeric substance (EPS) production at the experimental endpoint. EPS composition was characterized by measuring protein, lipid, and carbohydrate fractions within the total EPS pool.

Neither protein nor carbohydrate EPS differed significantly with diversity or nutrient treatment (**Figure 3AB**). However, lipid EPS was significantly associated with diversity (p = 0.02), with high-diversity nutrient-supplemented mesocosms containing the greatest lipid EPS concentration (0.157 ± 0.01 mg g⁻¹ soil), followed by high-diversity nutrient-deficient (0.08 ± 0.01 mg g⁻¹ soil), low-diversity nutrient-supplemented (0.05 ± 0.01 mg g⁻¹ soil), and low-diversity nutrient-deficient mesocosms (0.02 ± 0.01 mg g⁻¹ soil; **Figure 3C**).

**Figure 3.**
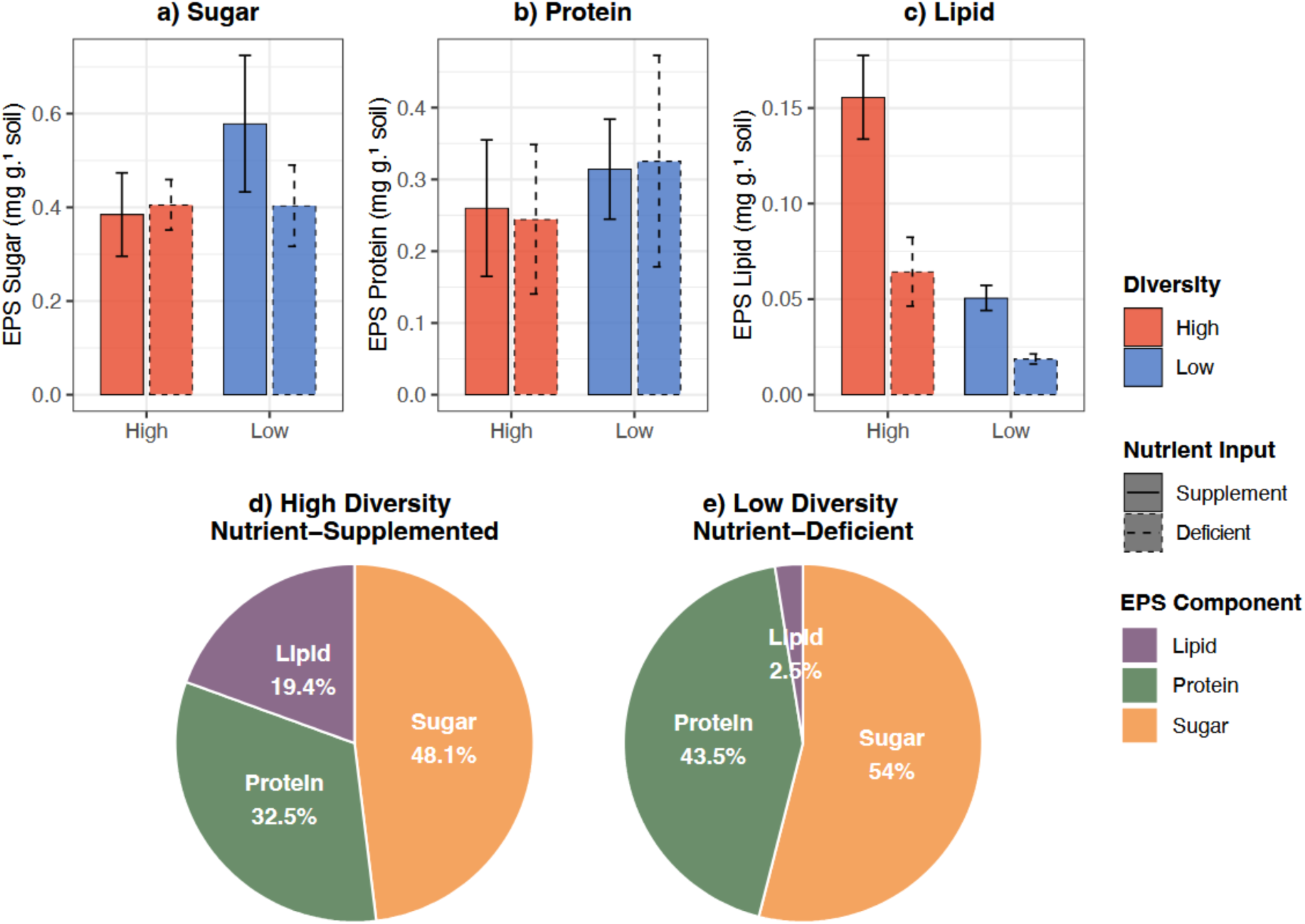
Extracellular polymeric substance (EPS) composition of soil mesocosms under contrasting diversity and nutrient treatments. Panels (a–c) show mean (± SE) concentrations of carbohydrate (a), protein (b), and lipid (c) EPS fractions (mg g⁻¹ soil) for high-diversity (red) and low-diversity (blue) mesocosms under nutrient-supplemented (solid bars) and nutrient-deficient (dashed outline bars) conditions. Panels (d) and (e) show the proportional composition of the total EPS pool for the high-diversity nutrient-supplemented (d) and low-diversity nutrient-deficient (e) treatments. n = 3 mesocosms per treatment combination.

Comparison of overall EPS composition between the high-diversity nutrient-supplemented and low-diversity nutrient-deficient treatments revealed the greatest compositional divergence occurred in the lipid fraction, which was 16.9% higher in the high-diversity nutrient-supplemented treatment (**Figures 3DE**). In contrast, protein and carbohydrate fractions were proportionally higher in the low-diversity nutrient-deficient treatment, with protein accounting for 11 percentage points more and carbohydrates 5.9 percentage points more of the total EPS pool. Together, these results suggest that EPS composition, rather than total EPS quantity, responds most strongly to differences in diversity and nutrient status, with lipid EPS in particular tracking the diversity effect observed in water retention.

### Microbiome within Mesocosms

Amplicon sequencing was performed to verify that microbial diversity manipulations were successfully maintained within the mesocosms across the experimental period. Sequencing was carried out on the inoculum and within mesocosms at mid-sampling and at the experimental endpoint.

Inoculum alpha diversity confirmed the intended diversity manipulation, with the high-diversity inoculum containing 610 observed ASVs compared to just 5 ASVs in the low-diversity inoculum. These patterns were maintained within the mesocosms throughout the experiment. High-diversity nutrient-supplemented mesocosms had a mean observed alpha diversity of 110 ASVs, high-diversity nutrient-deficient mesocosms averaged 500 ASVs, low-diversity nutrient-supplemented mesocosms averaged 30 ASVs, and low-diversity nutrient-deficient mesocosms averaged 50 ASVs (**Figure 4a**). The notably higher alpha diversity in nutrient-deficient mesocosms within the high-diversity treatment warrants further investigation and may reflect competitive release under nutrient limitation. Microbial community composition differed significantly between treatments, with treatment identity (diversity and nutrient input in combination) explaining 42% of the variation in community composition (PERMANOVA: F = 5.069, p < 0.001; **Figure 4b**). Across all treatments, microbial communities were predominantly composed of Firmicutes and Proteobacteria, though nutrient-supplemented treatments showed greater representation of *Actinobacteriota, Planctomycetota*, and *Verrucomicrobiota*. Clear clustering by diversity level was observed in the NMDS ordination, confirming that the diversity manipulation successfully and persistently altered microbial community structure within the mesocosms.

**Figure 4.**
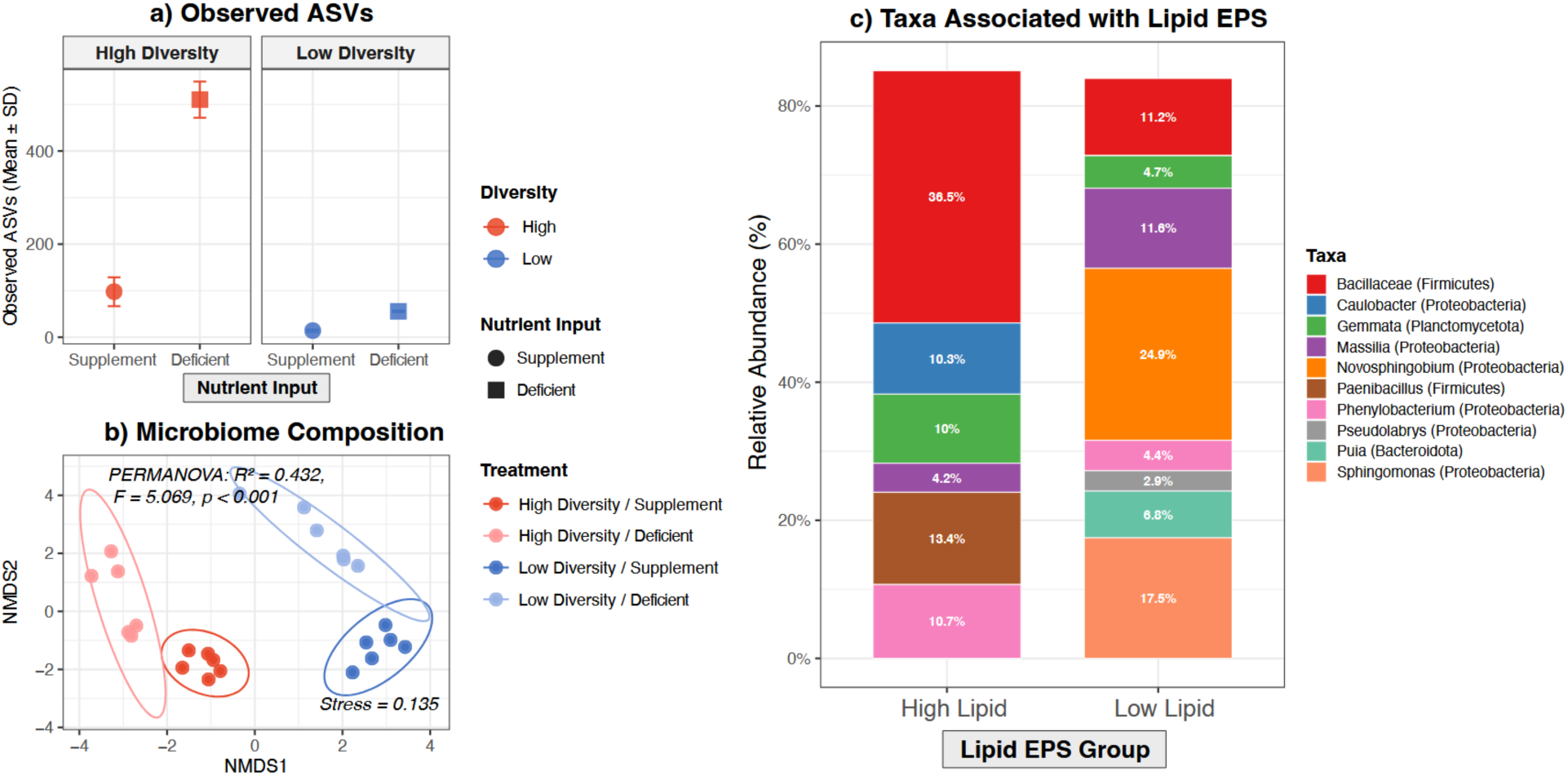
Microbial community characteristics of soil mesocosms. Panel (a) shows mean (± SD) observed ASV richness for high-diversity (red) and low-diversity (blue) mesocosms under nutrient-supplemented (circles) and nutrient-deficient (squares) conditions. Panel (b) shows non-metric multidimensional scaling (NMDS) ordination of microbial community composition based on Bray-Curtis dissimilarity, with 95% confidence ellipses shown per treatment group. PERMANOVA results and NMDS stress value are annotated within the panel. Panel (c) shows the relative abundance of taxa identified as significantly associated with lipid EPS concentration by DESeq2 analysis (padj < 0.05), comparing samples grouped by median lipid EPS into high and low lipid groups. Taxa are colored by identity; unlabeled slices represent taxa contributing less than 1% relative abundance.

Given that lipid EPS was identified as the primary EPS fraction responding to diversity and nutrient treatments (**Figure 3**), we investigated whether specific microbial taxa were associated with variation in lipid EPS concentration. PERMANOVA analysis revealed that lipid concentration was the only EPS component to explain significant variation in microbial community composition (PERMANOVA: F = 2.481, p = 0.009), accounting for 10% of compositional variation, while glucose and protein concentrations were non-significant. DESeq2 analysis identified 12 ASVs significantly associated with lipid EPS concentration (padj < 0.05; **Figure 4c**). Taxa positively associated with higher lipid EPS included *Bacillaceae, Caulobacter,* and *Paenibacillus*, while taxa associated with lower lipid EPS included Massilia, *Novosphingobium*, and *Sphingomonas*, suggesting distinct microbial functional groups may drive lipid EPS production and degradation within these systems.

### Integrating Hydrological, Microbial, and EPS Responses

To investigate relationships between EPS fractions and soil hydrological properties, we conducted regression analyses between sugar, protein, and lipid EPS concentrations against average and peak water loss and infiltration rates. Of all EPS fractions, only lipid EPS showed a significant relationship with hydrology, exhibiting a negative association with peak water loss rate (r = -0.705, p = 0.010; **Figure 5a**), while protein and sugar EPS showed no significant relationships with either water retention or infiltration (**Figure 5b–c**, p > 0.05).

**Figure 5.**
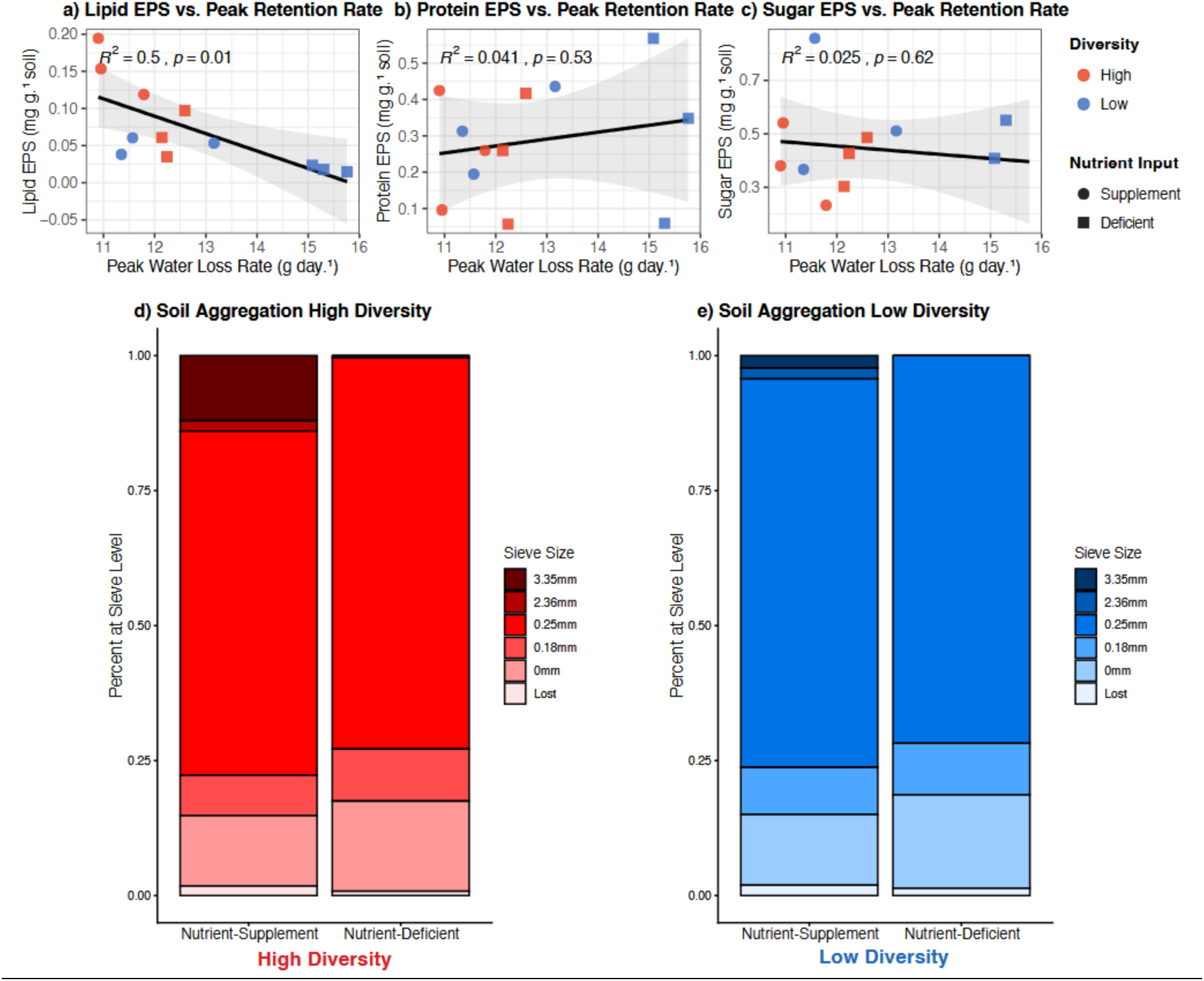
Relationship between extracellular polymeric substance (EPS) fractions and peak hydrological rates across soil mesocosms. Panels (a–c) show the relationship between peak water loss rate and lipid (a), protein (b), and sugar (c) EPS concentrations. Panels (d–f) show the relationship between peak infiltration rate and lipid (d), protein (e), and sugar (f) EPS concentrations. Peak water loss rate was calculated as the maximum single-day decline in water volume retained per mesocosm across the second rewetting cycle; peak infiltration rate was calculated as the maximum single-day increase in water volume infiltrated to the bottom cup. Points are colored by diversity treatment (red = high diversity, blue = low diversity) and shaped by nutrient treatment (circles = nutrient-supplemented, squares = nutrient-deficient). Regression lines represent a single pooled linear fit across all treatment combinations; shaded bands indicate 95% confidence intervals. Pearson correlation coefficients (R²) and associated p-values are annotated in each panel. n = 3 mesocosms per treatment combination.

To contextualize these findings within broader soil physical mechanisms, we examined soil aggregate size distributions and bulk soil properties. Aggregate size increased with both microbial diversity and nutrient availability, mirroring the pattern observed in water retention data (**Figure 2, 5d–e**). The proportion of large aggregates (> 3.35 mm) was greatest in the high-diversity nutrient-supplemented treatment (12.5%), declining substantially in the low-diversity nutrient-supplemented treatment (∼5%). Within high-diversity mesocosms, the large aggregate proportion declined further from 12.5% under nutrient supplementation to less than 2% under nutrient deficiency, with a similar reduction observed in low-diversity systems.

Bulk soil carbon and nitrogen measurements revealed that organic carbon via combustion was negatively associated with average water loss rate (r = -0.841, p = 0.001; **Figure 6**), while elevated inorganic nitrogen, both ammonium (r = 0.684, p = 0.014; **Figure 6**) and nitrate (r = 0.674, p = 0.016; **Figure 6**), was positively associated with faster average water loss rates, suggesting that nutrient availability is related to potentially altered soil hydrological properties beyond its direct effects on microbial community composition and EPS production. When considering peak rather than average rates, nitrate remained positively associated with peak water loss (r = 0.703, p = 0.011; **Figure 6**), while lipid EPS was negatively associated with peak water loss (r = -0.705, p = 0.010; **Figure 5a, 6**), indicating that mesocosms with greater lipid EPS concentrations experienced lower peak water loss events. Together these results suggest that lipid EPS and inorganic nitrogen availability may act in opposing directions on peak soil water loss, with lipid EPS buffering against extreme water loss events while elevated nitrate is associated with more rapid peak drainage.

**Figure 6.**
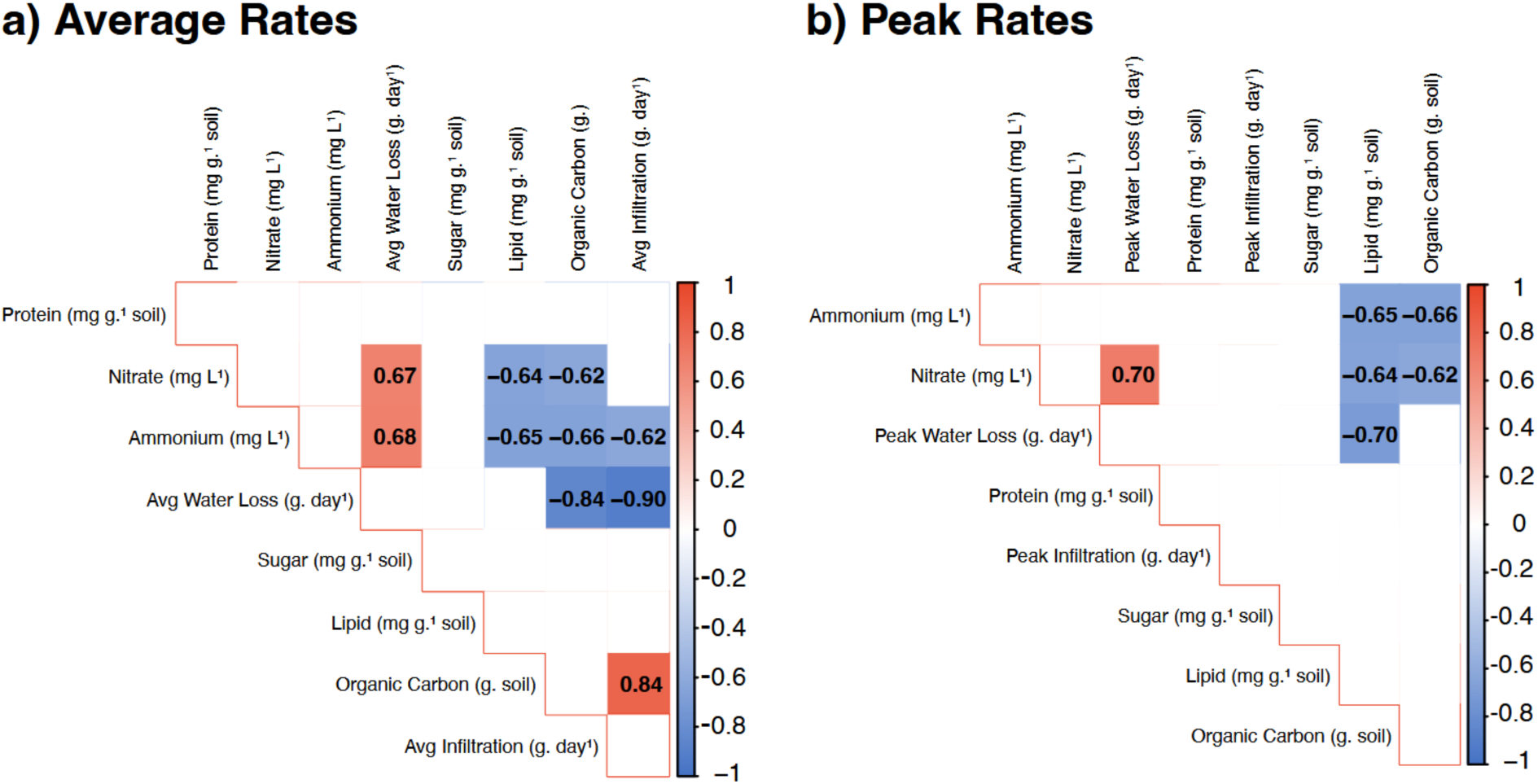
Pearson correlation matrices between EPS fractions, soil properties, and hydrological rates. Panel (a) shows correlations using average water loss and infiltration rates, while panel (b) shows correlations using peak single-day water loss and infiltration rates. Variables included are lipid, protein, and sugar EPS fractions (mg g⁻¹ soil), organic carbon (g), ammonium (mg L⁻¹), nitrate (mg L⁻¹), and average or peak hydrological rates (g day⁻¹). Correlation coefficients (r) are displayed within each cell; blank cells indicate non-significant correlations (p > 0.05).

## DISCUSSION

Soil microbial communities are multifunctional entities capable of shaping soil physical properties, including nutrient cycling, structural stability, and chemical composition (Philippot et al., 2024). In this study, we demonstrate that modulating microbial diversity in combination with nutrient availability drives measurable changes in soil hydrology within experimental mesocosms, with high-diversity nutrient-supplemented systems retaining significantly more water than low-diversity nutrient-deficient ones (**Figure 2**). We propose that this effect is mediated through microbial EPS production, specifically lipid EPS, which binds soil particles, promotes aggregate formation, and acts as a hydrophilic matrix capable of retaining water, establishing a mechanistic chain from microbial diversity through EPS production to soil physical structure and hydrological function.

While the influence of microbial diversity on soil ecosystem functions such as decomposition, nitrification, and denitrification is well established (Wagg et al., 2014; Delgado-Baquerizo et al., 2016), this study is among the first to demonstrate that community-level diversity contributes to soil hydrology through EPS-mediated mechanisms. Previous work has shown that individual high-EPS-producing isolates can alter soil hydraulic properties (Roberson and Firestone, 1992), and that gelatinous polysaccharides directly added to soils can modify water retention (Chenu, 1993), but the role of diversity per se has received little attention. Our results also demonstrate that nutrient addition further enhanced EPS production and water retention, though accompanied by a reduction in microbial diversity in some treatments, consistent with nutrient-diversity trade-offs reported by Zeng et al. (2016) and Yang et al. (2022). Critically, diversity retained its own independent effect on water retention even in nutrient-supplemented systems, suggesting these factors act through partially distinct mechanisms and that microbial diversity should be considered an important and underappreciated determinant of soil hydrological properties.

EPS production is a well-characterized microbial strategy for surviving fluctuating hydrological conditions, enabling cells to retain water during desiccation and resist osmotic stress (Costa et al. 2018). Within soil systems, EPS pools respond dynamically to land-use change, with greater concentrations observed in undisturbed compared to agronomically managed soils (Kidinda et al., 2023; Redmile-Gordon et al., 2020). However, much of this work has focused on carbohydrate-based EPS polymers. By characterizing major EPS fractions simultaneously, we found that lipid EPS was the most predictive of hydrological change; a result that was not anticipated given the conventional view of lipids as hydrophobic. Many microbial lipids are in fact amphiphilic, possessing both hydrophobic and hydrophilic domains, and may contribute to water retention through hydrophilic head groups while simultaneously stabilizing aggregates through hydrophobic interactions with mineral surfaces. Recent advances in soil lipidomics support this interpretation, Couvillion et al. (2023) demonstrated that the soil microbiome lipidome shifts substantially under desiccation, while Samrat et al. (2025) showed that lipid signatures across soil organisms were altered under warming and drought. Furthermore, recent work in streambed biofilms has shown that EPS concentrations increase under drought conditions (Romani et al., 2025), suggesting that EPS upregulation in response to drying may be a conserved microbial strategy across terrestrial and aquatic environments. Together, these findings implicate lipid remodeling and EPS accumulation as active microbial responses to water stress and suggest that the lipid EPS signal observed here may reflect a community-level strategy for modifying soil hydrological properties during drying.

The physical mechanism linking EPS to hydrology likely operates through aggregate formation. Organic matter is well established as a driver of aggregate stability (Tisdall and Oades, 1982; Six et al., 2004), and our findings extend this framework by identifying lipid EPS, modulated by diversity and nutrient availability, as a specific contributor to this process. The parallel patterns observed between aggregate size distribution, lipid EPS concentration, and water retention across treatments suggest these processes are tightly coupled, with microbial diversity acting as an upstream regulator of both EPS composition and the soil physical structure that determines hydrological behavior. Soil aggregation underpins a wide range of critical functions, including aeration, carbon sequestration, and erosion resistance (Bronick and Lal, 2005), and lipid EPS production represents a concrete biological target that could be monitored as a soil health metric. The specific taxa driving these effects provide further mechanistic insight. *Bacillaceae*, *Caulobacter*, and *Paenibacillus* were positively associated with lipid EPS, taxa well characterized as EPS producers. *Paenibacillus* and *Bacillaceae* are known to produce exopolysaccharides (Liyaskina et al., 2021; Marvasi et al., 2010), and *Caulobacter* is recognized for its holdfast EPS structures facilitating surface attachment (Sprecher et al., 2017). Different microbial species produce compositionally distinct EPS fractions (Flemming and Wingender, 2010), and the co-occurrence of multiple EPS-producing taxa in high-diversity mesocosms may generate synergistic interactions or functional redundancy that ensures EPS production is maintained even if individual taxa are lost (Louca et al., 2018). Conversely*, Massilia, Novosphingobium*, and *Sphingomonas* were negatively associated with lipid EPS, potentially acting as EPS degraders, certain species are known to produce enzymes that degrade heterospecific EPS pools as a competitive strategy (Flemming et al., 2016; Pontrelli et al., 2025). The balance between EPS-producing and EPS-degrading taxa may therefore be a key determinant of net EPS accumulation and its hydrological consequences.

These findings carry important applied implications. Targeted management of soil microbial diversity could represent a novel strategy for improving soil moisture retention in agricultural systems, particularly relevant given the well-documented decline in soil water availability driven by increasing drought frequency and irregular precipitation (Yao et al., 2025, Seo et al., 2025), and the growing dependence on irrigated agriculture placing pressure on global freshwater resources (Sauer et al., 2010). While the present experiment was conducted without plants, a critical next step will be determining whether EPS-mediated water retention benefits extend to plant-soil systems and whether these effects can buffer crops against drought stress, a question with direct relevance to food security under climate change.

Several limitations must be acknowledged. The controlled mesocosm setting, while enabling precise manipulation, may not fully capture field soil complexity including plant roots, soil fauna, and spatial heterogeneity. The small sample size limits statistical power, and replication across soil types and field conditions will be essential. Before these findings can inform practical applications, three questions must be addressed: whether EPS-mediated hydrological effects are reproducible in intact field soils; what the specific biological mechanisms and genetic pathways underlying microbial EPS production are; and how microbial diversity can be practically manipulated to improve water use efficiency at scale.

Together, these findings demonstrate that microbial diversity and nutrient availability interact to shape soil hydrological properties through EPS-mediated mechanisms, with lipid EPS emerging as a key functional link between community composition and water retention capacity.

As soil degradation and drought frequency increase globally, managing microbial diversity represents an underexplored but promising solution for enhancing soil hydrological resilience.

## Acknowledgements

We thank the University of Arizona College of Agriculture, Life and Environmental Sciences, and the School of Plant Sciences for supporting this work. We gratefully acknowledge to all the members of the Favela Lab that assisted with this work: A. DeRusha, L. Bowdren, J. Marcos, C. Hale, P. Nitz, C. Curry, F. Gomez., M. Banach, and G. Steen. Furthermore, we want to acknowledge the staff of the University of Arizona Genetics Core (UAGC), namely J. Galina-Mehlam with assistance. We further would like to acknowledge UBRP director J. Cubeta for excellent leadership of the program.

## Author Contributions

Writing—original draft: Y.K and A.F. Conceptualization: A.F., Investigation: Y.K., and A.F. Writing—review and editing: Y.K., M.A, H.B. E.H. A.S and A.F. Methodology: Y.K. and A.F. Resources: A.F. Funding acquisition: Y.K. and A.F., Data curation: Y.K. and A.F., Supervision: A.F.

